# Ectopic activation of the polar body extrusion pathway triggers cell fragmentation in preimplantation embryos

**DOI:** 10.1101/2022.12.22.521568

**Authors:** Diane Pelzer, Ludmilla de Plater, Peta Bradbury, Adrien Eichmuller, Anne Bourdais, Guillaume Halet, Jean-Léon Maître

**Affiliations:** Institut Curie, PSL Research University, CNRS UMR3215, INSERM U934, 75005 Paris, France; Institut de Génétique et Développement de Rennes, Université de Rennes, CNRS UMR 6290, Rennes, France

## Abstract

Cell fragmentation occurs during physiological processes, such as apoptosis, migration, or germ cell development. Fragmentation is also commonly observed during preimplantation development of human embryos and is associated with poor implantation prognosis during Assisted Reproductive Technology (ART) procedures. Despite its biological and clinical relevance, the mechanisms leading to cell fragmentation are unclear. Light sheet microscopy imaging of mouse embryos reveals that compromised spindle anchoring, due to Myo1c knockout or dynein inhibition, leads to fragmentation. We further show that defective spindle anchoring brings DNA in close proximity to the cell cortex, which, in stark contrast to previous reports in mitotic cells, locally triggers actomyosin contractility and pinches off cell fragments. The activation of actomyosin contractility by DNA in preimplantation embryos is reminiscent of the signals mediated by small GTPases throughout polar body extrusion (PBE) during meiosis. By interfering with the signals driving PBE, we find that this meiotic signaling pathway remains active during cleavage stages and is both required and sufficient to trigger fragmentation. Together, we find that fragmentation happens in mitosis after ectopic activation of actomyosin contractility by signals emanating from DNA, similar to those observed during meiosis. Our study uncovers the mechanisms underlying fragmentation in preimplantation embryos and, more generally, offers insight into the regulation of mitosis during the maternal-zygotic transition.

## Main text

Cell fragmentation can lead to the complete disintegration of a cell, aiding clearance during processes such as apoptosis (Atkin-Smith and Poon, 2017). Additionally, cell fragmentation can also be partial and contribute to cell and tissue maturation including during germ cell or gonad formation in c. elegans embryos (Abdu et al., 2016; Lee et al., 2019), or the deposition of signaling migrasomes during zebrafish gastrulation (Jiang et al., 2019). In other instances, the function and/or benefit of cell fragmentation remains unclear. Interestingly, fragmentation is commonly observed during the preimplantation development of human embryos and is correlated with a poor implantation outcome in Assisted Reproductive Technologies (ART). In human embryos, fragmentation is often associated with chromosome mis-segregation and the formation of micronuclei, negatively affecting further embryonic development (Alikani, 1999; Alikani, 2007; Fujimoto et al., 2011). Importantly, fragmentation in human embryos occurs naturally (Pereda and Croxatto, 1978) and independently of apoptosis (Hardy, 1999). While cell fragmentation is frequently observed in human embryos and has been reported in rhesus monkey embryos (Daughtry et al., 2019; Hurst et al., 1978), it is rarely observed in other mammalian species including mice (Chavez et al., 2012) - the typical model used to study human preimplantation development. Therefore, identifying mouse mutants that recapitulate human embryo fragmentation would serve as a useful model to study fragmentation during mammalian preimplantation development.

While studying preimplantation morphogenesis, we serendipitously found that loss of unconventional myosin-Ic (Myo1c) causes partial fragmentation in mouse embryos. Myo1c, one of the two Myo1 paralogs expressed during early preimplantation development (Deng et al., 2014) (Fig S1a), tethers actin filaments to phosphatidylinositol 4,5-biphosphate-containing membranes (Lebreton et al., 2018; McIntosh and Ostap, 2016) found at the plasma membrane of preimplantation embryos (Halet et al., 2008). We find that zygotic KO of Myo1c (Myo1cKO) after CRISPR/Cas9 microinjection causes embryos to form numerous large fragments (Fig 1a). In Myo1cKO embryos, fragments were 5-10 times more numerous than in control embryos and sized twice as large (Fig 1a–c, Fig S1b). Fragments in Myo1cKO morula rarely contained DNA and were found at the apical surface of the embryo (Fig S1c–d), similar to that described for both human and rhesus monkey embryos (Daughtry et al., 2019; Hardy, 1999; Van Blerkom et al., 2001). Fragments that contained DNA were found in the blastocoel of both control and Myo1cKO blastocysts (Fig S1b–d) and are the likely result of complete cell fragmentation triggered by programmed cell death within the inner cell mass (Plusa et al., 2008). Therefore, Myo1cKO mouse embryos form fragments sharing the characteristic location and content of those found in human embryos.

**Figure 1:**
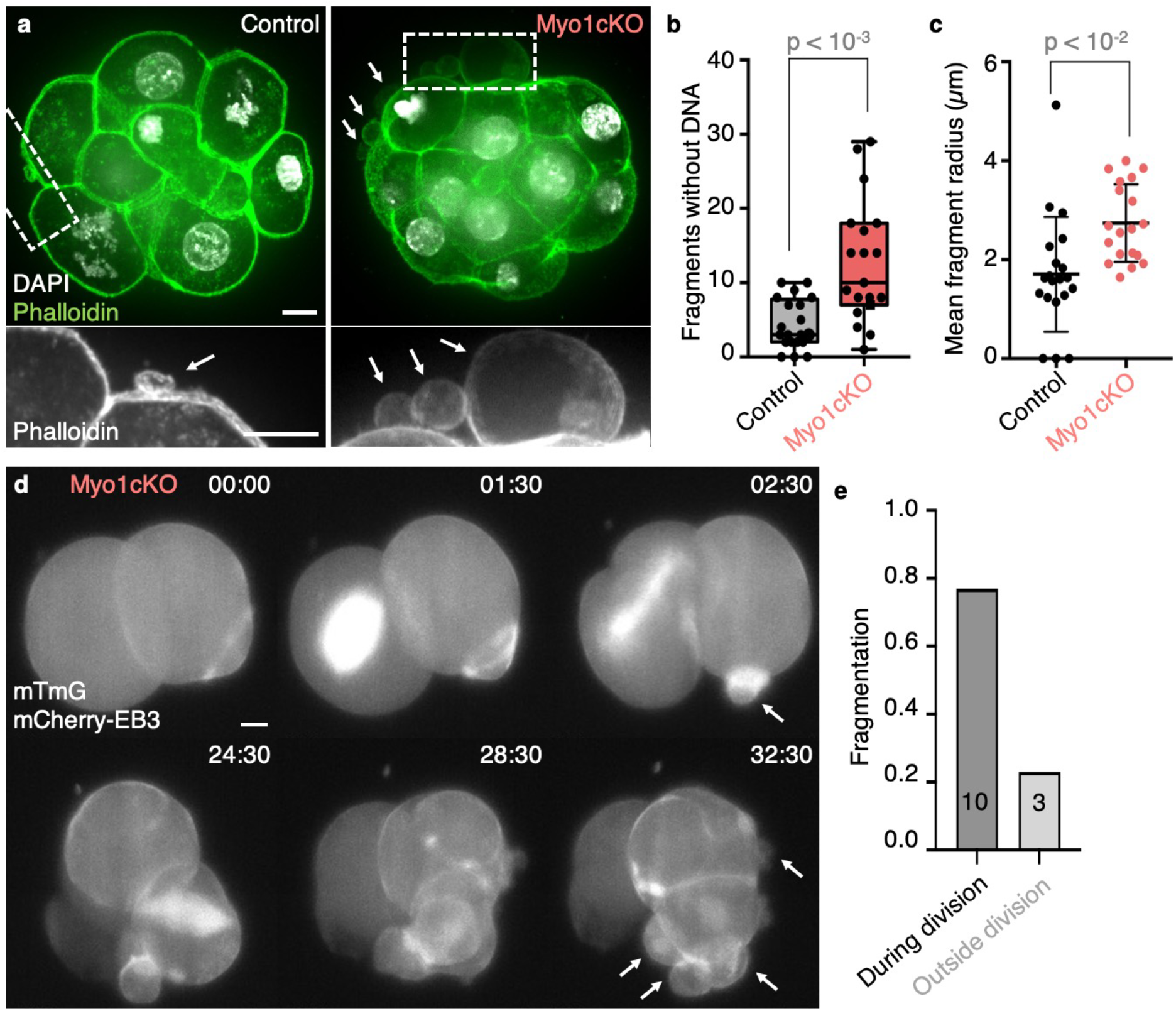
Myo1cKO embryos form fragments (**a**) Representative image of a Control (left) and Myo1cKO (right) embryo. Top: max projections of embryos stained with Phalloidin (green) and DAPI (gray). Bottom: magnifications at dashed rectangles of Phalloidin (grey) with white arrows pointing at fragments. (**b-c**) Number of fragments without DNA (**b**) per embryo and mean radius (**c**) in Control (grey) and Myo1cKO (salmon) embryos (left, Control n = 20, Myo1cKO n = 20). (**d**) Representative images of a time-lapse of Myo1cKO embryo undergoing fragmentation outside of division (top) and during division (bottom). Max projections of embryos expressing mTmG (grey) and mCherry-EB3 (grey) imaged with light sheet microscopy are shown. White arrows point at the forming fragments. (**e**) Ratio of fragmentation events during and outside of division (n = 13 embryos). Mann-Whitney test *p* values are indicated. Scale bars, 10 *μ*m.

Fragmentation in human embryos tends to occur during mitosis and in clusters (Van Blerkom et al., 2001). To investigate precisely when fragmentation takes place in Myo1c mutants, we used light-sheet microscopy of Myo1cKO embryos expressing a plasma membrane marker (mTmG) to visualize fragmentation and mCherry-EB3 to identify mitotic cells by the presence of the mitotic spindle. This revealed that few fragments formed during interphase, and most fragmentation events coincided with the presence of the mitotic spindle (Fig 1d–e, Movie 2). In addition, we observed that fragments formed both individually and in clusters (Fig 1d, Movie 2). Therefore, like human embryos, Myo1cKO mouse embryos form fragments during mitosis and in clusters.

After division, fragmenting blastomeres were to some extent able to divide and develop further (Movie 1). Early developmental processes such as compaction and apico-basal polarization appeared to occur normally in Myo1cKO embryos (Fig S1f–h, Movie 1), which also initially contained a similar number of cells as control embryos (Fig S1b, e). However, Myo1cKO embryos eventually failed to form normal blastocysts. At the blastocyst stage, Myo1cKO embryos exhibited reduced cell number compared to control embryos (Fig S1b, e) and showed impaired lineage specification and lumen formation (Fig S1i–m). Therefore, as in human embryos (Hardy, 1999; Van Blerkom et al., 2001), cell fragmentation in Myo1cKO mouse embryos is not necessarily linked to cell death but is associated with poor developmental potential to the blastocyst stage.

In summary of the phenotypic analysis of zygotic Myo1cKO embryos, we found that large apical DNA-free fragments form during mitosis without immediately affecting cell viability but compromising preimplantation development, as reported in human embryos. Thus, using Myo1cKO mouse embryos, which form fragments with the attributes of those typically found in human embryos, we set out to dissect the mechanisms of fragmentation in preimplantation embryos.

While imaging the mitotic spindle in Myo1cKO embryos, we noted several differences with the spindle of control embryos (Fig 2a, Movie 3). In control embryos, the spindle visualized with mCherry-EB3 appeared and persisted for ~50 min whereas, in Myo1c mutants, the spindle lingered for ~100 min (Fig 2b). Furthermore, we noted that the spindle rotated and bent extensively in Myo1cKO embryos when compared to control ones (Fig 2c–d). Therefore, cell fragmentation in Myo1cKO embryos is associated with poor mitotic spindle anchoring, as described in vitro following Myo1c knockdown (Mangon et al., 2021). To test if faulty spindle anchoring would cause fragmentation independently of Myo1c KO, we weakened spindle anchoring using Ciliobrevin D to inhibit dynein, the protein responsible for tethering astral microtubules to the cell cortex (Firestone et al., 2012; Kotak et al., 2012). Embryos treated with Ciliobrevin D prior to cell division showed enhanced persistence and rotation of the mitotic spindle when compared to DMSO treated embryos (Fig S2), confirming that inhibition of dynein impaired spindle anchoring. Strikingly, this treatment phenocopied Myo1c knockout and promoted cell fragmentation (Fig 2e–f). Together, we find that Myo1c or dynein dysfunctions are two possible disruptions causing both spindle anchoring defects and cell fragmentation, raising the possibility that spindle anchoring defects could be involved in fragmentation.

**Figure 2:**
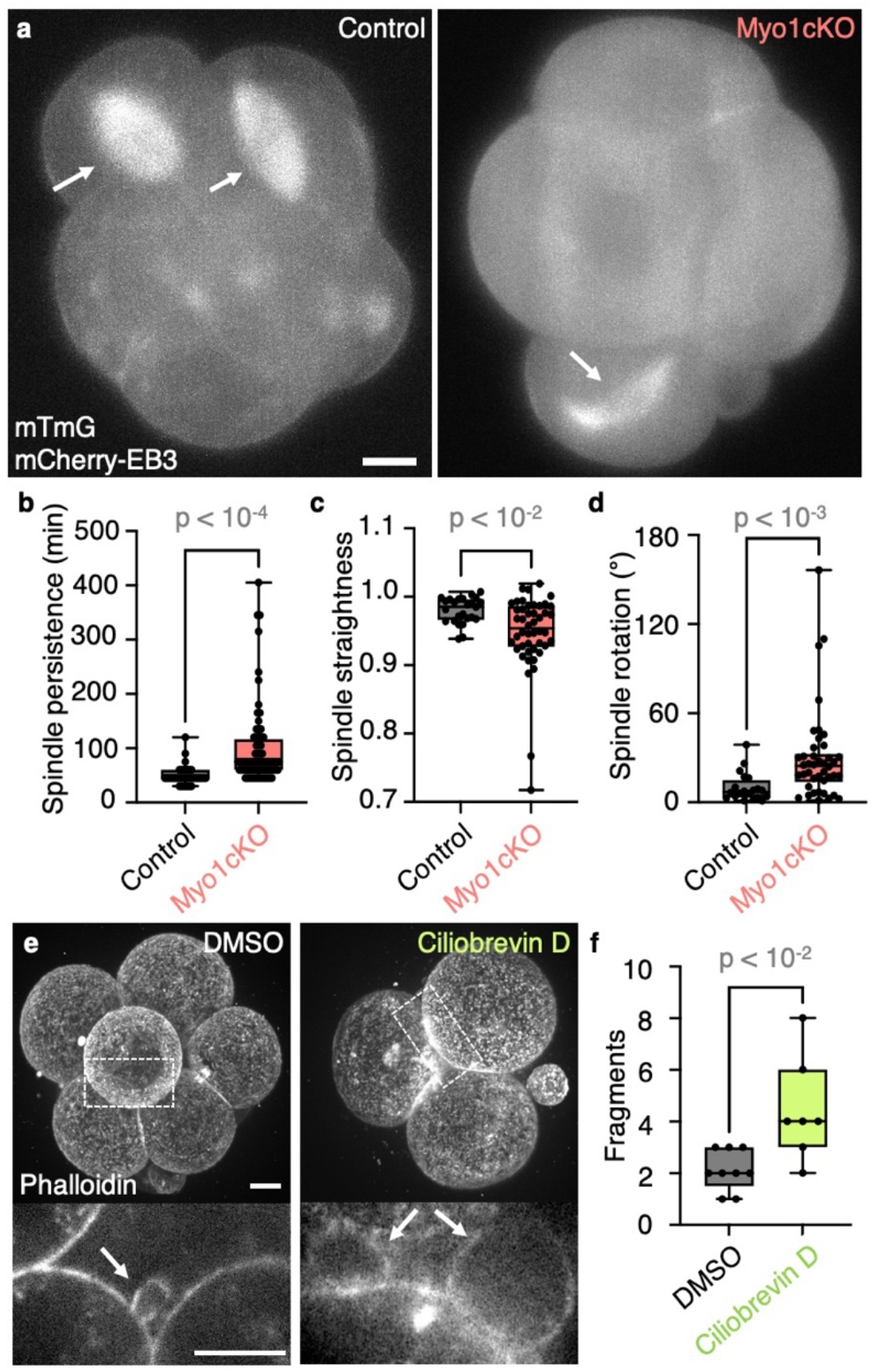
Defective spindle anchoring leads to fragmentation (**a**) Representative images of Control (left) and Myo1cKO (right) embryos expressing mTmG and mCherry-EB3 (grey) shown as max projections. White arrows point at the mitotic spindle. (**b-d**) Spindle persistence (**b**), straightness (**c**) and rotation (**d**) in Control (n = 6) and Myo1cKO (n = 9) embryos. Persistence measures the time from appearance to disassembly of the spindle. Straightness reports the pole to pole distance of the spindle divided by its continuous length. Rotation describes the angle between the initial orientation of the spindle pole and its maximal deviation in 2D. (**e**) Representative images of embryos treated with DMSO (left) and 37.5 μM Ciliobrevin D (right), stained with Phalloidin. Top: Maximum projection. Bottom: magnifications of a single plane at dashed rectangles of Phalloidin with white arrows pointing at fragments. (**f**) Number of fragments without DNA per embryo in DMSO (grey) and Ciliobrevin D (honeydew) embryos at E2.5 (left, Control n = 9, Myo1cKO n = 7). Mann-Whitney test *p* values are indicated. Scale bars, 10 *μ*m.

Early divisions of human preimplantation embryos frequently show defective spindles, causing chromosome misalignments and segregation errors (Cavazza et al., 2021; Ford et al., 2020; Kalatova et al., 2015). Since the spindle in Myo1cKO embryos is defective, we suspected that chromosomes may be incorrectly positioned during mitosis. Indeed, labeling chromosomes with SiRDNA or H2B-mCherry revealed that chromosomes in Myo1cKO embryos often come in close proximity with the cell cortex instead of remaining at the center of dividing cells (Fig 3a, Movie 4). Previous studies reported that, during mitosis, chromosomes induce the relaxation of the actomyosin cortex when in close proximity to the pole of dividing cells (Kiyomitsu and Cheeseman, 2013; Ramkumar et al., 2021; Rodrigues et al., 2015; Sedzinski et al., 2011).

**Figure 3:**
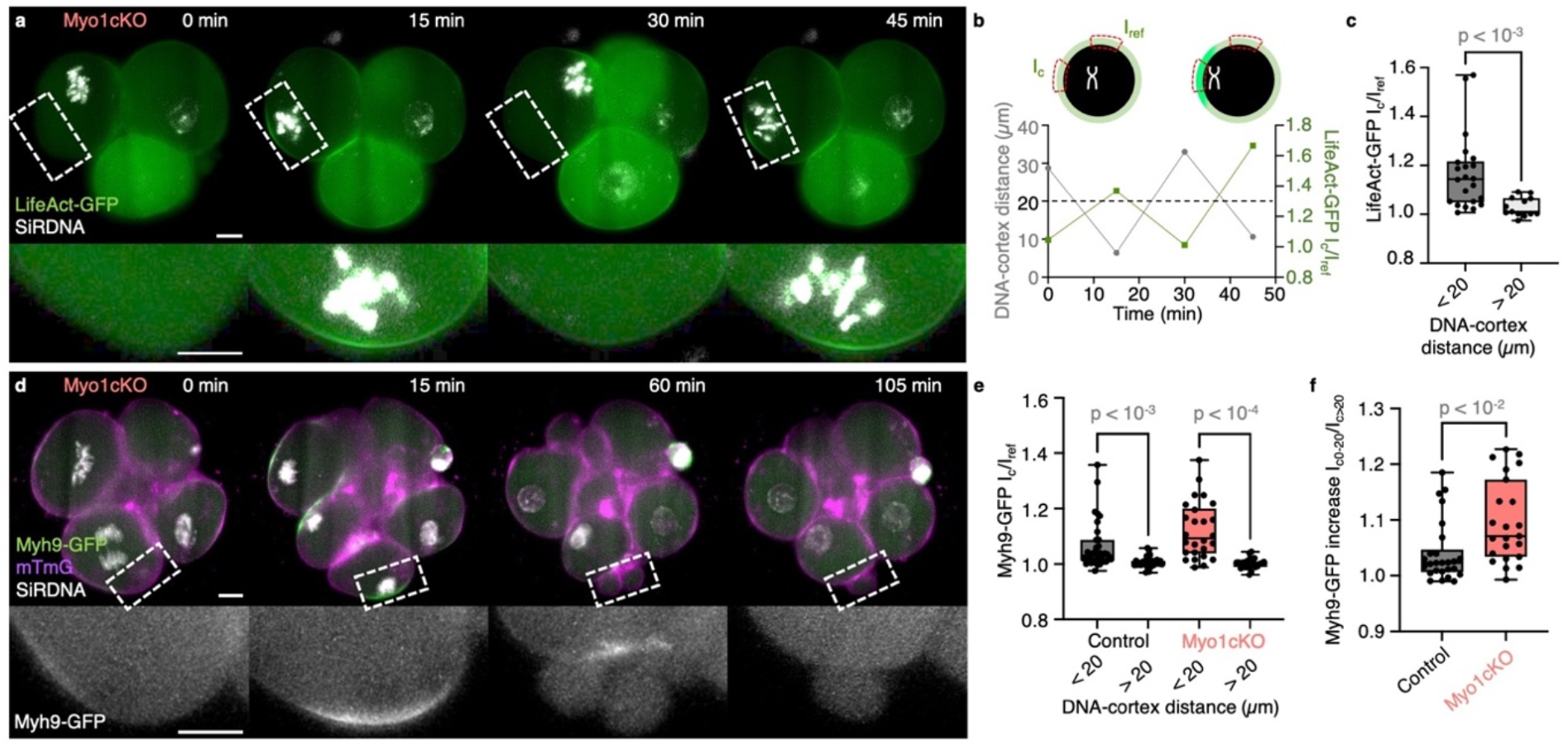
Actomyosin cap formation in proximity of DNA (**a**) Top: Representative images of a timelapse of a Myo1cKO embryo labeled with LifeAct-GFP (green) and SiRDNA (grey) with one blastomere undergoing mitosis shown as max projections. Bottom: magnifications of dashed rectangles. (**b**) In Myo1cKO embryo shown in (**a**) LifeAct-GFP intensities at the cortex region of eventual contact with DNA (I_c_) are normalized to the intensity at a reference region (I_ref_). LifeAct-GFP I_c_/I_ref_ values (green) are plotted over time. The DNA-cortex distance (gray) between the region of eventual contact and the center of the DNA signal is also measured. A threshold distance of 20 μm is indicated with a dashed line. (**c**) LifeAct-GFP Ic/Iref values at DNA-cortex distances < 20 *μ*m and > 20 *μ*m (Myo1cKO embryos n = 13, cells n = 21). (**d**) Top: Representative images of a time-lapse of a Myo1cKO embryo expressing Myh9-GFP (green) and mTmG (magenta) while labelled with SiRDNA (grey) undergoing fragmentation during mitosis shown as max projections. Bottom: magnifications of Myh9-GFP (grey) at dashed rectangles showing cortical accumulation (middle) followed by fragmentation (right). (**e**) Control (grey) and Myo1cKO (salmon) embryos Myh9-GFP I_c_/I_ref_ values at DNA-cortex distances < 20 *μ*m and > 20 *μ*m (Control embryos n = 10, cells n = 35; Myo1cKO embryos n = 9, cells n = 26) (**f**) Comparison of the increase in Myh9-GFP intensity in Control and Myo1cKO embryos when DNA is in close proximity to the cortex. The ratio I_c0-20_/I_c>20_ is calculated from Myh9-GFP intensities at the contact region when DNA-cortex distances are < 20 *μ*m (I_c0-20_) divided by intensities when distances are > 20 *μ*m (I_c>20_) (Control embryos n = 10, cells n = 33; Myo1cKO n = 9, cells n = 26). Mann-Whitney test (**c**, **f**) or Kruskal-Wallis and pairwise Mann-Whitney tests (**e**) *p* values are indicated. Scale bars, 10 *μ*m.

In stark contrast, we noted that when chromosomes neared the cortex of mitotic cells of Myo1cKO embryos, actin accumulated at the cortex, as seen using the filamentous actin reporter LifeAct (Fig 3a–c, Movie 4-5). Remarkably, at the poles, non-muscle myosin IIA, visualized using Myh9-GFP, reached levels comparable to, or higher than that present at the cleavage furrow (Movie 5). Therefore, not all cells relax their poles during mitosis.

Measuring the local levels of LifeAct-GFP or Myh9-GFP in Myo1cKO embryos revealed increased intensity once DNA came within 20 *μ*m of the cortex (Fig 3b–e). Importantly, the location of Myh9-GFP accumulation corresponded to the site of fragment formation (Fig 3d, Movie 4-5). Careful observation of the formation of fragments suggested that Myh9-GFP formed a ring-like structure that ‘pinched off’ a piece of the cell (Movie 6). Importantly, when DNA came within close proximity to the cortex, Myh9-GFP accumulation could be detected in both control and Myo1cKO embryos (Fig 3e). However, Myh9-GFP accumulation in control embryos was less pronounced than in Myo1cKO ones (Fig 3f), suggesting that the signaling cascade that induces cortical Myh9 accumulation is more stimulated and/or more responsive in Myo1cKO embryos than in the control ones. Together, our findings indicate that spindle anchoring defects bring chromosomes closer to the cortex for an extended duration (Fig 2) and coincides with ectopic activation of contractility, causing a small portion of the cell to pinch off, thereby generating a fragment (Fig 3).

The aforementioned series of events described in mitotic cells, strikingly resembles the extrusion of the polar body during meiosis (Uraji et al., 2018). In mouse oocytes, the meiotic spindle must migrate towards the cell periphery forcing chromosomes within close proximity of the actomyosin cortex. The meiotic spindles and chromosomes eventually triggers a signaling cascade that assembles an actin cap surrounded by a non-muscle myosin-II ring to pinch off the polar body. Strikingly, the assembly of the actomyosin cap extruding the polar body is induced when chromatin is within 20 *μ*m of the cortex (Deng et al., 2007), a distance similar to that observed prior to fragment formation during cleavage stages (Fig 3b, c, e). Two distinct signals coordinate polar body extrusion (PBE). The first originates from the chromosomes and involves, among other effectors, the small GTPase, Cdc42 to promote actin polymerization (Deng et al., 2007; Dumont et al., 2007). The second emanates from the central spindle, which organizes the contractile cytokinetic ring, via, among other effectors, the Rho Guanine Exchange Factor (GEF), Ect2 (Dehapiot et al., 2021). Importantly, the PBE signaling cascade is responsible for both encapsulating half of the maternal genome into the polar body, and also for repelling the remaining maternal chromosomes away from the cortex (Dehapiot et al., 2021; Wang et al., 2020). In fact, following the completion of meiosis, paternal chromosomes that near the cortex also induce actin caps and can form transient buds in zygotes (Mori et al., 2021; Simerly et al., 1998). Therefore, the signaling pathway driving PBE remains active hours after the completion of meiosis and it is unclear when this meiotic pathway is terminated.

To test if the PBE pathway is active in cleavage stage embryos and involved in cell fragmentation, we investigated the two signals involved in PBE - those that originate from the chromosomes (via Cdc42) and from the spindle (via Ect2). We monitored Cdc42 activity using a probe that exploits the Cdc42-GTP-binding region of myotonic dystrophy kinase-related Cdc42-binding kinase (MRCK) (Bourdais et al., 2022). We observed that the active Cdc42 probe accumulated at the cortex specifically when chromosomes were within 20 *μ*m (Fig 4a–b, Movie 7), suggesting that, like during meiosis, the PBE pathway is active during mitosis. Importantly, Cdc42 was activated in both control and Myo1cKO embryos without necessarily triggering fragment formation, similar to when paternal DNA induces the formation of actin caps without being extruded (Mori et al., 2021; Simerly et al., 1998). To next determine whether Cdc42 activation is responsible for the chromosome-induced actin accumulation in Myo1cKO embryos, we used a dominant negative Cdc42 (DNCdc42). As described previously, when chromosomes came within 20 *μ*m of the cortex of Myo1cKO embryos, an accumulation of actin was observed (Fig 3a–c) however, this was abolished by the expression of DNCdc42 (Fig 4c–d, Movie 8). Therefore, fragmentation of preimplantation embryos is driven by Cdc42-mediated actin polymerization following DNA nearing the cortex. In addition, we investigated signals originating from the mitotic spindle by increasing the levels of Ect2. Expression of Ect2-GFP induced the formation of numerous large fragments, while expression of GFP alone did not (Fig 4e–f). Therefore, increased signaling from Ect2 is sufficient to induce fragment formation. Taken together, we find that core elements of the PBE pathway are active, necessary, and sufficient to induce fragmentation in preimplantation embryos.

**Figure 4:**
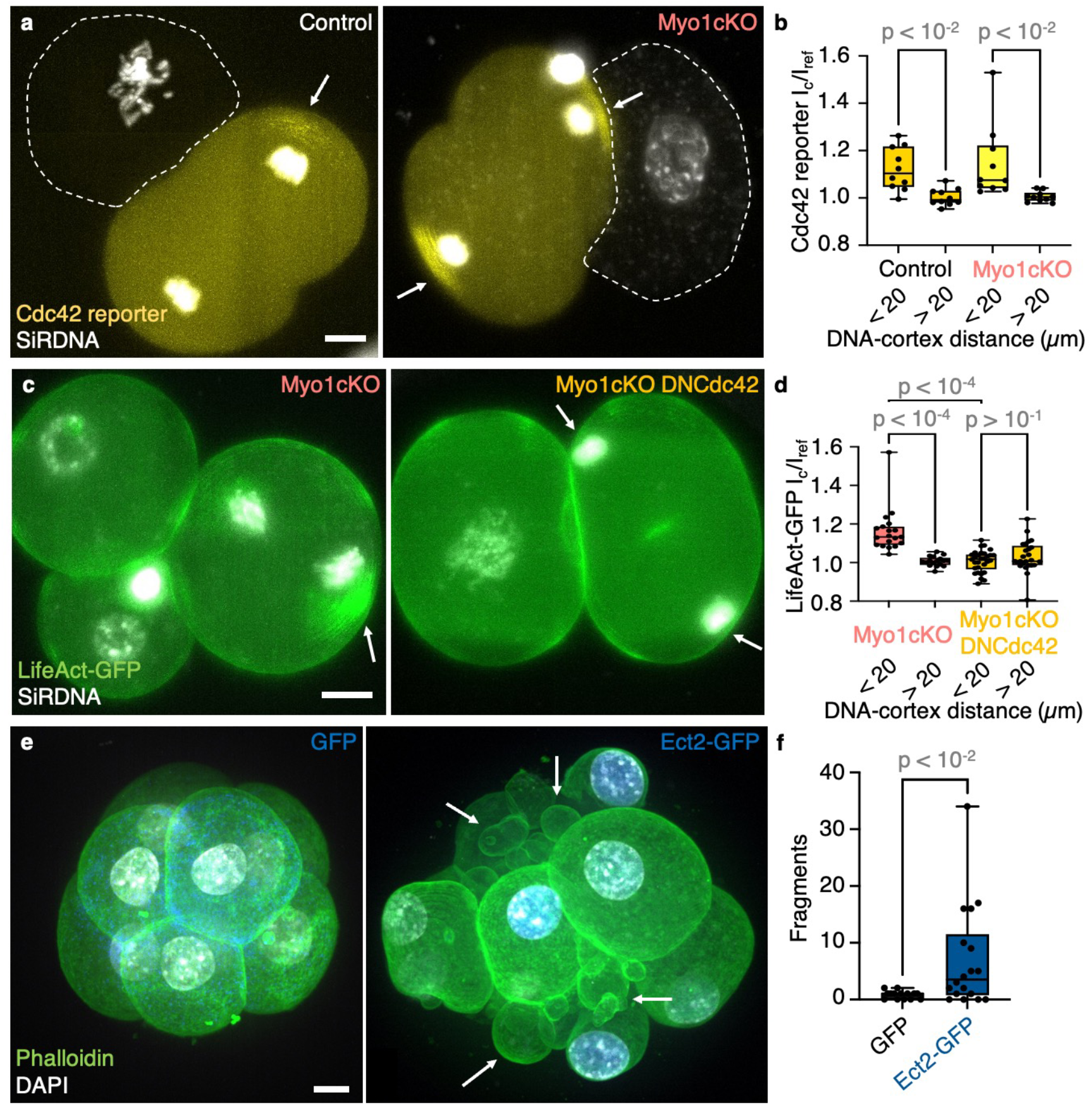
The polar body extrusion (PBE) pathway is necessary and sufficient to induce fragmentation during cleavage stages. (**a**) Representative images of Control (left) and Myo1cKO (right) embryos labelled with SiRDNA (grey) and expressing a reporter of Cdc42 activity (yellow) in one blastomere, which is undergoing mitosis, shown as max projections. The second blastomere is outlined with a dashed line. White arrows point at the accumulation of active Cdc42 at the cortex in close proximity to DNA. (**b**) The ratio of active Cdc42 I_c_/I_ref_ was compared between < 20 *μ*m and > 20 *μ*m DNA-cortex distances for Control (dark yellow) and Myo1cKO (yellow) embryos (Control embryos n = 10; Myo1cKO embryos n = 10). (**c**) Representative images of Myo1cKO embryos expressing LifeAct-GFP (green) alone (left) or together with dominant negative Cdc42 (DNCdc42, right) labelled with SiRDNA (grey) shown as max projections. White arrows point at the cortex in close proximity to DNA. (**d**) LifeAct-GFP I_c_/I_ref_ values at DNA-cortex distances < 20 *μ*m and > 20 *μ*m for Myo1cKO (salmon) and Myo1cKO + DNCdc42 (yellow) embryos (Myo1cKO embryos n = 8, cells n = 19; Myo1cKO + DNCdc42 embryos n = 12, cells n = 30). (**e**) Representative images of a GFP (left) and Ect2-GFP (right) expressing embryo stained with Phalloidin (green) and DAPI (grey) shown as max projections. White arrows indicate fragments. (**f**) Number of fragments without DNA per embryo in GFP (grey) and Ect2-GFP (blue) embryos (GFP n = 14; Ect2-GFP n = 18). Kruskal-Wallis and pairwise Mann-Whitney tests (**b**, **d**) or Mann-Whitney test (**f**) *p* values are indicated. Scale bars, 10 *μ*m.

Our experiments show that ectopic signals originating from the chromosomes or from the spindle can lead to fragmentation. To dissect the specific mechanisms of action of both signals, we measured actomyosin levels upon DNA nearing the cortex as a readout of PBE pathway activation. In Myo1cKO embryos, Myh9-GFP recruitment was increased compared to control embryos (Fig 3f), which could be explained by the longer persistence of the mitotic spindle dragging the chromosomes throughout the cell (Fig 2b, Movie 3 and 4). Therefore, when the mitotic spindle is poorly anchored, the PBE pathway appears to be hyper-activated compared to control embryos. In Ect2-GFP expressing embryos, actin levels increased when DNA neared the cortex as observed in control embryos (Fig S3a–b, Movie 9). However, in contrast to Myo1c KO, Ect2-GFP expression did not enhance actin recruitment compared to control embryos (Fig S3b). Therefore, hyperactivation of the PBE pathway may not be the cause of fragmentation in Ect2-GFP expressing embryos.

Since Ect2 activates Rho, a master regulator of actomyosin contractility (Matthews et al., 2012), expression of Ect2-GFP could raise the basal contractility and lower the threshold for fragmentation upon normal activation of the PBE pathway. To assess contractility, we used micropipette aspiration to measure the surface tension of embryos expressing Ect2-GFP (Maître et al., 2015). Surface tension of blastomeres expressing Ect2-GFP was increased when compared to GFP-expressing cells (Fig S3c). In contrast, Myo1cKO embryos displayed surface tensions that were identical to control embryos (Fig S1g–h). Together, these measurements indicate that, even when the PBE pathway is ectopically activated to normal levels, fragmentation can occur when contractility is abnormally high. In summary, we observe fragmentation either after hyper-activation of the PBE pathway in cells with normal contractility or after ectopic activation of the PBE pathway in cells with hyper contractility.

Taken together, our experiments delineate a mechanism to explain cell fragmentation (Fig 5), a process that is detrimental to human embryonic development and human fertility. We find that fragments form by ectopic contraction of the actomyosin cortex (Fig 3) upon activation of a meiotic pathway that is unexpectedly still operative during mitosis (Fig 4). Specifically, we find that the PBE pathway is engaged, as during meiosis, when chromosomes come near the cortex (Fig 3–4). During mitosis, ectopic activation of the PBE pathway is more pronounced when the spindle suffers from poor anchoring (Fig 2). Since human embryos show frequent issues with spindle and chromosome capture (Cavazza et al., 2021; Ford et al., 2020; Kalatova et al., 2015), we propose that fragmentation may result from hyper-activation of the PBE pathway due to problems during chromosome separation. Importantly, ectopic activation of the PBE pathway occurs in control embryos without causing fragmentation (Fig 3–4). However, in embryos with increased basal contractility and surface tension, the same ectopic activation of the PBE pathway triggers fragmentation (Fig 4 and Fig S3). Interestingly, in a recent study, we measured the surface tension of human embryos and reported tensions 5-10 times higher than for mouse embryos (Firmin et al., 2022). Thus, with unstable spindles and high contractility, human embryos cumulate two features that could synergistically promote fragmentation (Fig 5). Future studies focusing on reducing contractility and/or PBE pathway activation in human embryos may therefore be key to improve human fertility.

**Figure 5:**
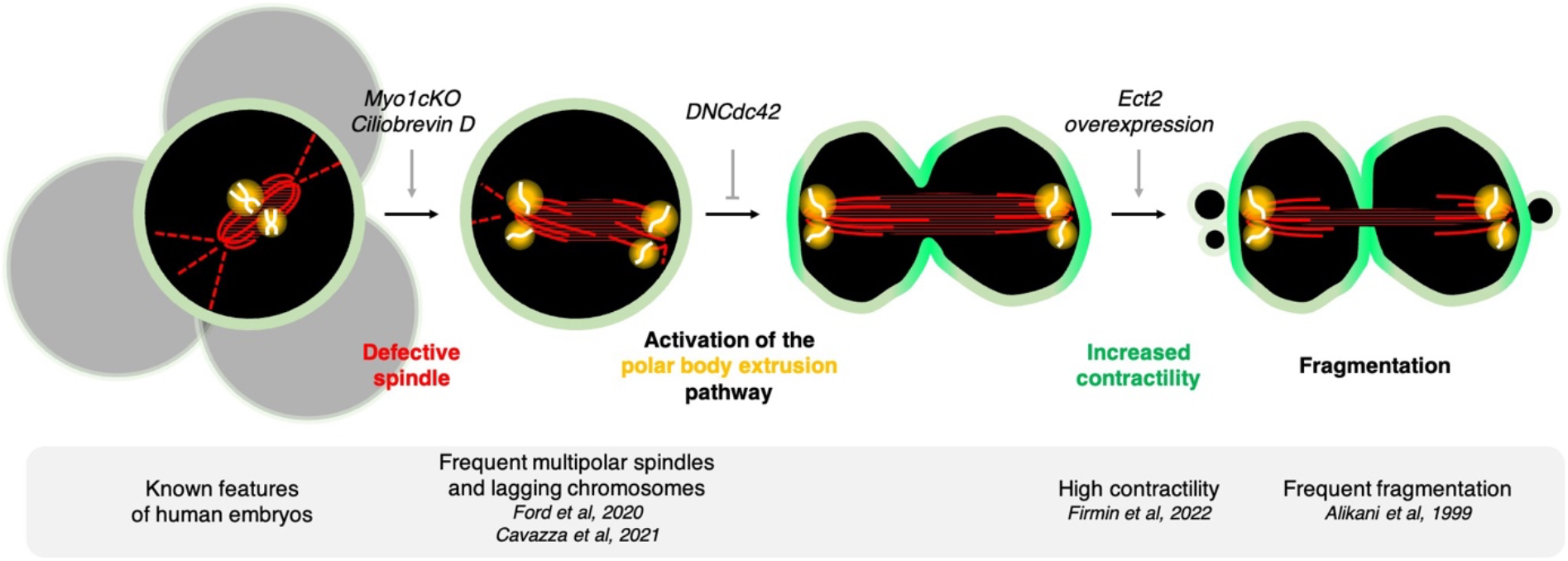
Schematic diagram of the steps leading to cell fragmentation in preimplantation embryos. Spindle in red, DNA in white, signals from the DNA in yellow and actomyosin in green.

Finally, we uncover novel features of mitotic regulation. We find that not all mitotic cells relax the acto-myosin cortex when chromosomes come near the cortex, contrary to what has been described so far in somatic cells (Kiyomitsu and Cheeseman, 2013; Ramkumar et al., 2021; Rodrigues et al., 2015; Sedzinski et al., 2011). Specifically, we show in cleavage stage embryos, when DNA comes within 20 *μ*m of the cortex, it locally activates Cdc42 and triggers actomyosin contraction (Fig 3–4). We propose that this is due to the persistent activity of the PBE pathway 3 days after completion of meiosis (Fig 4), revealing an unexpected prolongation of maternal processes in the embryo. There may be in fact little pressure to terminate the PBE pathway upon completion of meiosis. Indeed, cleavage stage blastomeres are rather large cells, thereby reducing the risk that a well anchored and centered mitotic spindle would bring chromosomes near the cortex. We recently found that, as a result of remodeling of the actomyosin cortex, surface tension decreases concomitantly with cells decreasing in size during cleavage stage embryos (Özgüç et al., 2022). This progressive mechanical maternal-zygotic transition may reduce the consequences of the ectopic activation of the PBE pathway and prevent cell fragmentation.

Whether cortical softening occurs in human embryos remains unknown and our findings highlight that much remains to be understood about how embryos transition both molecularly and mechanically from meiotic to mitotic division.

## Acknowledgements

We thank the imaging platform of the Genetics and Developmental Biology unit at the Institut Curie (PICT-IBiSA@BDD) for their outstanding support; the animal facility of the Institut Curie for their invaluable help. We thank Marie-Émilie Terret, Marie-Hélène Verlhac, Kristine Schauer, Jean-Baptiste Brault and Alexandre Baffet for sharing constructs and discussions. We thank Marie-Hélène Verlhac, Yohanns Bellaïche and members of the Maître lab for critical reading of the manuscript.

Research in the lab of J.-L.M. is supported by the Institut Curie, the Centre National de la Recherche Scientifique (CNRS), the Institut National de la Santé Et de la Recherche Médicale (INSERM), and is funded by grants from the ATIP-Avenir program, the Fondation Schlumberger pour l’Éducation et la Recherche via the Fondation pour la Recherche Médicale, the European Research Council Starting Grant ERC-2017-StG 757557, the Agence Nationale de la Recherche (ANR-21-CE13-0010-02), the European Molecular Biology Organization Young Investigator program (EMBO YIP), the INSERM transversal program Human Development Cell Atlas (HuDeCA), Paris Sciences Lettres (PSL) QLife (17-CONV-0005) grant and Labex DEEP (ANR-11-LABX-0044) which are part of the IDEX PSL (ANR-10-IDEX-0001-02).

## Author contributions

D.P., L.P., P.B. and A.E. performed experiments and prepared data for analyses. A.B. and G.H. provided constructs and insights into the polar body extrusion pathway. D.P. and J.-L.M. designed the project, analyzed the data, and wrote the manuscript. J.-L.M. acquired funding.

## Methods

### Embryo work

#### Recovery and culture

All animal work is performed in the animal facility at the Institut Curie, with permission by the institutional veterinarian overseeing the operation (APAFIS #11054-2017082914226001). The animal facilities are operated according to international animal welfare rules.

Embryos are isolated from superovulated female mice mated with male mice. Superovulation of female mice is induced by intraperitoneal injection of 5 international units (IU) pregnant mare’s serum gonadotropin (PMSG, Ceva, Syncro-part), followed by intraperitoneal injection of 5 IU human chorionic gonadotropin (hCG, MSD Animal Health, Chorulon) 44-48 hours later.

Embryos are recovered at E0.5 in 37°C FHM (LifeGlobal, ZEHP-050 or Millipore, MR-122-D) by dissecting, from the oviduct, the ampula, from which embryos are cleared with a brief (5-10 s) exposure to 37°C hyaluronidase (Sigma, H4272).

Embryos are recovered at E1.5 by flushing oviducts from plugged females with 37°C FHM using a modified syringe (Acufirm, 1400 LL 23). Embryos are handled using an aspirator tube (Sigma, A5177-5EA) equipped with a glass pipette pulled from glass micropipettes (Blaubrand intraMark or Warner Instruments).

Embryos are placed in KSOM (LifeGlobal, ZEKS-050 or Millipore, MR-107-D) or FHM supplemented with 0.1 % BSA (Sigma, A3311) in 10 *μ*L droplets covered in mineral oil (Sigma, M8410 or Acros Organics) unless stated otherwise. Embryos are cultured in an incubator with a humidified atmosphere supplemented with 5% CO2 at 37°C.

To remove the Zona Pellucida (ZP) for tension measurments, embryos are incubated for 45-60 s in pronase (Sigma, P8811).

For imaging, embryos are placed in Viventis Microscopy LS1 Live light-sheet chambers or 5 glass-bottom dishes (MatTek).

Only embryos surviving the experiments were analyzed. Survival is assessed by continuation of cell division as normal when embryos are placed in optimal culture conditions.

#### Mouse lines

Mice are used from 5 weeks old on. (C57BL/6xC3H) F1 hybrid strain is used for wild-type (WT). To visualize plasma membranes, mTmG (Gt(ROSA)26Sor^tm4(ACTB-tdTomato,-EGFP)Luo^) is used (Muzumdar et al., 2007). To visualize LifeAct-GFP (Tg(CAG–EGFP)#Rows) is used (Riedl et al., 2010). To visualize H2B-mCherry R26-H2B-mCherry CDB0239K is used (Abe et al., 2011). To generate embryos with a maternal Myh9-GFP allele, Myh9^tm8.1RSad^ females were mated with WT males (Zhang et al., 2012).

#### Chemical reagents and treatments

Ciliobrevin D (Merck, 250401) 5 mM dimethyl sulfoxide (DMSO) stock was diluted to 37.5 *μ*M in KSOM. Late 2-cell-stage embryos were cultured in 37.5 *μ*M Ciliobrevin D for 1 day. Only those embryos that continued to divide were considered.

For live DNA staining, embryos were incubated in KSOM containing SiRDNA for 30 min prior to imaging.

#### Plasmids and mRNA preparation

The following plasmids were used: mCherry-EB3-7 (Addgene 55037), Ect2-GFP (Dehapiot et al., 2021), DNCdc42 (encoding Cdc42 with T17N swap) (Dehapiot et al., 2013), Cdc42 reporter (Bourdais et al., 2022), LifeAct-GFP, LifeAct-RFP (Riedl et al., 2008). For mRNA generation of mCherry-EB3 and Ect2-GFP, the template was generated by amplification of the fragments encoding mCherry-EB3 or Ect2-GFP respectively followed by SV40 PolyA with a forward (fwd) primer containing a T7site at its 5’ end. The GFP template was generated by amplification of the GFP sequence from the pCS2-GFP-Fmnl3 with a fwd primer containing the SP6 site. The remaining plasmids were linearized by restriction enzyme digestion. The linear fragments were used as template for mRNA transcription using the mMessage mMachine T7, T3 or SP6 kit (Invitrogen, AM1340, AM1344, AM1348) according to manufacturer’s instructions followed by resuspension in RNase free-water.

#### gRNA design and generation

To target Myo1c a set of two gRNAs for each gene targeting close to the start codon are designed using the web tool CHOPCHOP (Labun et al., 2019), namely TGACGGGGTTCGAGTGACCATGG and TTGACTGCCCGAGACCGGGTAGG. For RNA transcription, the gRNA target sequences are cloned into pX458 (Addgene, 48138) as previously described (Cong et al., 2013). Briefly, following digestion of the plasmid with BbsI, two annealed oligos encoding the gRNA target region, containing the 5’ overhang AAAC and the 3’ overhang CAAAAT are phosphorylated and cloned into pX458. The gRNA and its backbone are amplified by PCR with a fwd primer containing a T7 site. Following PCR purification RNA is transcribed using the MEGAshortscript T7 transcription kit (Invitrogen, AM1354). The RNA is then purified by using the MEGAclear kit (Invitrogen, AM1908).

#### Genotyping

Following imaging individual embryos are collected for genomic DNA extraction. Single embryos are placed into a genomic DNA extraction buffer (10 mM Tris pH 8.5, 50 mM KCl, 0.01% gelatin, Proteinase K). Subsequently, two nested PCRs were performed amplifying the region surrounding the predicted mutation site using the following primer pairs for Myo1c: fwd: CCTGTAACACGTAGCTGGGA rev: GGGGTTTGGATGGGGTCAT for the first PCR and fwd: GCCTGTGTCCAAAGTGTCCC rev: CCGTGCTCATGGCACTCAC for the second PCR.

WT and mutants were determined by gel electrophoresis. Both homozygous and heterozygous embryos are considered as Myo1cKO embryos and are pooled together in our analyses.

#### Microinjection

Glass capillaries (Harvard Apparatus glass capillaries with 780 *μ*m inner diameter) are pulled using a needle puller and microforge to build a holding pipette and an injection needle. The resulting injection needles are filled with RNA and or protein solution diluted in injection buffer (5mM Tris-HCl pH = 7.4, 0.1mM EDTA) to the following concentrations: mCherry-EB3 100 ng/*μ*L, Ect2-GFP, LifeAct-GFP, LifeAct-RFP and Cdc42T17N 200 ng/*μ*L and Cdc42 reporter 150 ng/*μ*L. To knock out Myo1c, zygotes were injected with 300 ng/*μ*L Cas9 protein (IDT, 1081058) and 80 ng/*μ*L gRNA1 and gRNA2 each diluted in injection buffer. The filled needle is positioned on a micromanipulator (Narishige MMO-4) and connected to a positive pressure pump (Eppendorf FemtoJet 4i). Embryos are placed in FHM drops covered with mineral oil under Leica TL Led microscope. Zygote or 2 cell-embryos were injected while holding with holding pipette connected to a Micropump CellTram Oil.

#### Micropipette aspiration

As described previously (Guevorkian and Maître, 2017; Maître et al., 2015), a microforged micropipette coupled to a microfluidic pump (Fluigent, MFCS EZ) is used to measure the surface tension of embryos. In brief, micropipettes of radii 8-16 *μ*m are used to apply step-wise increasing pressures on the cell surface until reaching a deformation, which has the radius of the micropipette (*R_p_*). At steady-state, the surface tension *γ* of the cell is calculated from the Young-Laplace’s law applied between the cell and the micropipette: *γ* = *P_c_* / 2 (1/*R_p_* - 1/*R_c_*), where *Pc* is the critical pressure used to deform the cell of radius of curvature *R_c_*.

Measurements of individual blastomeres from the same embryo are averaged and plotted as such.

#### Immunostaining

Embryos are fixed in 2% PFA (Euromedex, 2000-C) for 10 min at 37°C, washed in PBS and permeabilized in 0.01% Triton X-100 (Euromedex, T8787) in PBS (PBT) at room temperature before being placed in blocking solution (PBT with 3% BSA) at 4°C for 2-4 h. Primary antibodies are applied in blocking solution at 4°C overnight. After washes in PBT at room temperature, embryos are incubated with secondary antibodies and phalloidin in blocking solution at room temperature for 1 h. Embryos are washed in PBT and imaged in PBS-BSA immediately after.

| Primary antibodies | Dilution | Provider | RRID |
| --- | --- | --- | --- |
| Cdx2 | 1:200 | Abcam ab157524 | AB_2721036 |
| Sox2 | 1:100 | Abcam ab97959 | AB_2341193 |
| Pard6b | 1:50 | SantaCruz 166405 |  |
| Aqp3 | 1:500 | Novusbio NBP2-33872 |  |

| Secondary antibodies and dyes | Dilution | Provider | RRID |
| --- | --- | --- | --- |
| Alexa Fluor Plus 488 anti-mouse | 1:200 | Invitrogen, A32723 | AB_2633275 |
| Alexa Fluor 546 anti-mouse | 1:200 | Invitrogen, A11003 | AB_2534071 |
| Alexa Fluor Plus 488 anti-rabbit | 1:200 | Invitrogen, A32731 | AB_2633280 |
| Alexa Fluor Plus 546 anti-rabbit | 1:200 | Invitrogen, A11010 | AB_2534077 |
| Alexa Fluor 633 phalloidin | 1:200 | Invitrogen, A22284 |  |
| 4',6-diamidino-2-phenylindole (DAPI) | 1:1000 | Invitrogen, D1306 | AB_2629482 |
| SiRDNA | 1:1000 | Spirochrome |  |

#### Microscopy

Live imaging is performed using a Viventis Microscopy LS1 Live light-sheet microscope. Fluorescence excitation was achieved with a dual illumination scanned Gaussian beam light sheet of ~2.2 *μ*m full width at half maximum using 488 and 561 nm lasers. Signal was collected with a Nikon CFI75 Apo LWD 25x/1.1 objective and through 525/50-25 or 561/25 band pass filter onto an Andor Zyla 4.2 Plus sCMOS camera. The microscope is equipped with an incubation chamber to keep the sample at 37°C and supply the atmosphere with 5% CO 2.

Surface tension measurements are performed on a Leica DMI6000 B inverted microscope equipped with a 40x/0.8 DRY HC PL APO Ph2 (11506383) objective and Retina R3 camera and 0.7x lens in front of the camera. The microscope is equipped with an incubation chamber to keep the sample at 37°C and supply the atmosphere with 5% CO_2_.

Stained embryos are imaged on a Zeiss LSM900 Inverted Laser Scanning Confocal Microscope with Airyscan detector. Excitation is achieved using a 488 nm laser line through a 63x/1.4 OIL DICII PL APO objective. Emission is collected through a 525/50 band pass filter onto an airyscan photomultiplier (PMT) allowing to increase the resolution up to a factor 1.7.

### Bioinformatic analysis

Mouse and human single cell RNA sequencing data were extracted from (Deng et al., 2014) and (Yan et al., 2013), analyzed as in (De Iaco et al., 2017). In brief, single cell RNAseq datasets of human and mouse embryos (GSE36552, GSE45719) were downloaded and analyzed using the online platform Galaxy (usegalaxy.org). The reads were aligned to the reference genome using TopHat (Galaxy Version 2.1.1 (Kim et al., 2013)) and read counts generated with htseq-count (Galaxy Version 0.9.1 (Anders et al., 2015)). Normalized counts were determined with limma-voom (Galaxy Version 3.38.3 + galaxy3 (Law et al., 2014)).

### Data analysis

#### Image analysis

##### Fragment characterization

Using FIJI (Schindelin et al., 2012) to visualize individual confocal slices, fragments are identified as small spherical structures bounded by actin or membrane.

To measure fragment size, the confocal slice showing the largest fragment diameter is used and the contour of the membrane manually drawn.

The polar body, characterized by its presence from the zygote stage and intense DNA signal (much brighter than cells), was excluded from the analysis.

##### Cortical intensity measurements

FIJI is used to measure cortical intensity and distance to DNA. A segmented line is used to define the size and the location of the LifeAct or Myh9-GFP cap. Starting at the first division timepoint the intensity is measured at the segmented line, with the length of the eventually formed cap, and the distance between the center of the DNA mass and the center of the segmented line. The intensity of another segmented line drawn with the same length on a different part of the cell-medium interface where no cortical fluctuations are observed is used to normalize the intensity.

##### Spindle analysis

Spindle persistence was calculated by measuring the time between the appearance and disappearance of the spindle. Spindle straightness was measured using FIJI by dividing the length of a straight line connecting spindle poles divided by the length of a broken line drawn along the spindle. For spindle rotation, the angle between the initial direction of one spindle pole and its maximal change of direction was measured.

#### Statistics

Data are plotted using GraphPad Prism. Statistical tests are performed using GraphPad Prism. Statistical significance is considered when *p* < 10^-2^.

The sample size was not predetermined and simply results from the repetition of experiments. No sample that survived the experiment, as assessed by the continuation of cell divisions, was excluded. No randomization method was used. The investigators were not blinded during experiments.

### Data availability

The raw microscopy data, regions of interests (ROI) and analyses will be made available on a public repository.

---

**Figure S1:**
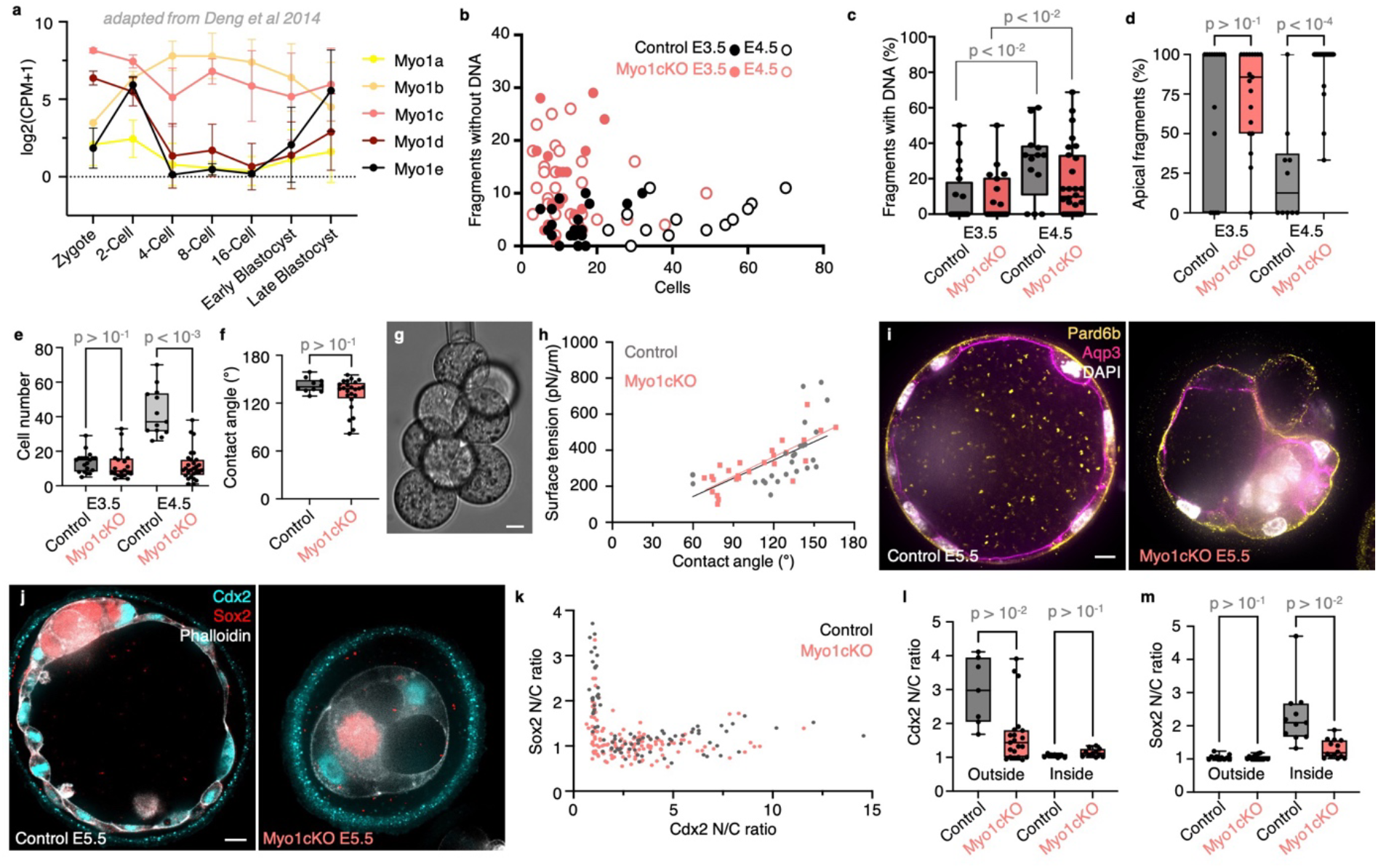
Characterization of Myo1cKO embryos development. (**a**) Mouse single cell RNA sequencing analysis adapted from Deng et al.(Deng et al., 2014), showing the expression levels of members for the Myosin 1 family at the zygote, 2-, 4-, 8-, 16-cell, early and late blastocyst stages. Data show mean ± SD. (**b-e**) Number of fragments without DNA as a function of the number of cells (**b**), percentage of fragments containing DNA per embryo (**c**), percentage of fragments located at the embryo apical surface (**d**), cell number per embryo (**e**) in Control (grey) and Myo1cKO (salmon) embryos at E3.5 (Control n = 20, Myo1cKO n = 20) and E4.5 (Control n = 13, Myo1cKO n = 28). (**f**) Mean contact angle at maximal compaction (Control n = 11, Myo1cKO n = 39). (**g**) Representative image of surface tension measurement. (**h**) Surface tension of control (grey, n = 6 embryos, 27 contacts) and Myo1cKO (salmon, n = 6 embryos, 53 contacts) blastomeres as a function of their contact angle. Representative image of tension measurement on an 8-cell stage embryo. (**i**) Representative images of Control (left) and Myo1cKO (right) embryos at E5.5 stained for Pard6b (yellow), Aqp3 (magenta) and DAPI (grey). (**j**) Representative images of Control (left) and Myo1cKO (right) embryos at E5.5 stained for Cdx2 (cyan) Sox2 (red) and Phalloidin (grey). (**k**) Sox2 nuclear to cytoplasmic (N/C) ratio as a function of Cdx2 N/C ratio of individual cells in Control (grey, n = 7) and Myo1cKO (salmon, n = 11) embryos. (**l**) Cdx2 N/C ratio of cells located at the surface (left) or inside (right) the embryo in Control (n = 8) and Myo1cKO (n = 21) embryos (**m**) Sox2 N/C ratio of cells located at the surface (left) or inside (right) the embryo in Control (n = 11) and Myo1cKO (n = 16) embryos. Mann-Whitney test (**f**) or Kruskal-Wallis test (**c, d, e, l, m**) *p* values are indicated. Scale bars, 10 *μ*m.

**Figure S2:**
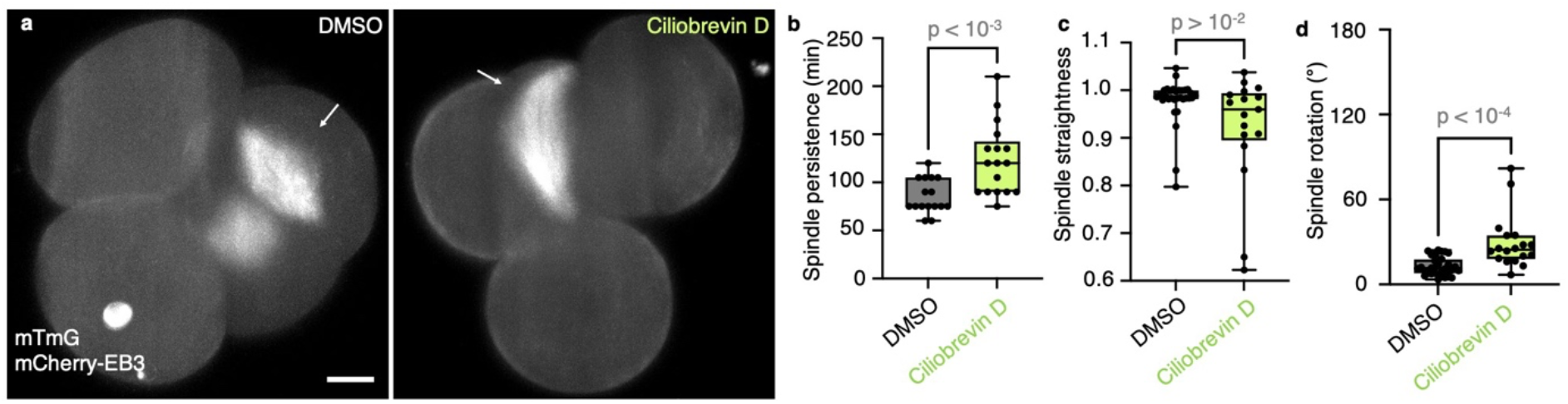
Mitotic spindle movement in Ciliobrevin D treated embryos. (**a**) Representative images of DMSO (left) and Ciliobrevin D (right) treated embryos expressing mTmG and mCherry-EB3 (grey) shown as max projections. White arrows indicate the mitotic spindle. Scale bar, 10 *μ*m. (**b-d**) Spindle persistence (**b**), straightness (**c**) and rotation (**d**) in DMSO (n = 11) and Ciliobrevin D (n = 11) treated embryos. Persistence measures the time from appearance to disassembly of the spindle. Straightness reports the pole to pole distance of the spindle divided by its continuous length. Rotation describes the angle between the initial orientation of the spindle pole and its maximal deviation in 2D. Mann-Whitney test *p* values are indicated.

**Figure S3:**
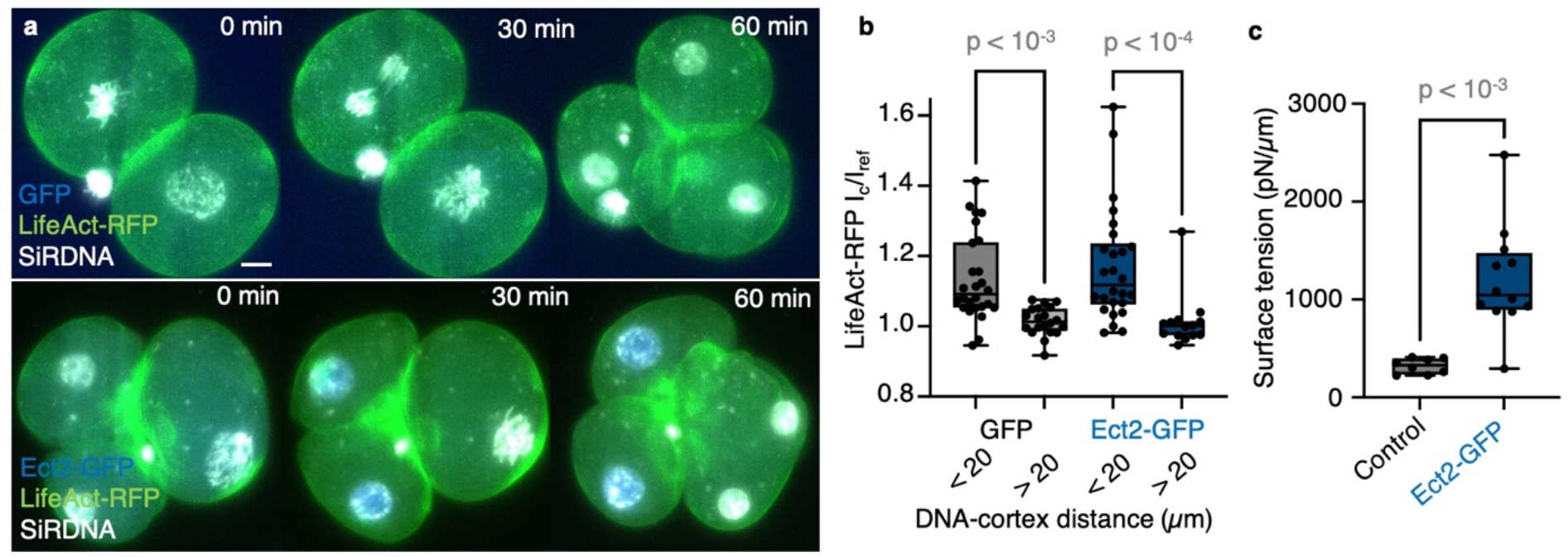
Characterisation of Ect2-GFP expressing embryos. (**a**) Representative images of GFP (top, blue) and Ect2-GFP (bottom, blue) expressing embryos during mitosis labeled with LifeAct-RFP (green) and SiRDNA (white) shown as max projections. Scale bar, 10 *μ*m. (**b**) LifeAct-RFP I_c_/I_ref_ ratio compared between DNA-cortex distances < 20 *μ*m and > 20 *μ*m for GFP (grey) and Ect2-GFP (blue) embryos (GFP embryos n = 7, cells n = 26; Ect2-GFP embryos n = 9, cells n = 26). (**c**) Surface tension of 4-cell stage embryos in Control (grey) and Ect2-GFP (blue) embryos (Control n = 8; Ect2-GFP n = 12). Kruskal-Wallis and pairwise Mann-Whitney tests (**b**) or unpaired Student’s *t* test (**c**) *p* values are indicated.

## Movie legends

**Movie 1: Myo1cKO embryos fragment and continue preimplantation development.**

Time lapse of a Myo1cKO embryo expressing mTmG (grey) undergoing fragmentation during the 3^rd^ and 4^th^ cleavages while proceeding with preimplantation development (compaction and polarization are visible). Images were taken every 15 min. Scale bar, 10 *μ*m. White arrows indicate fragments.

**Movie 2: Fragment formation during and outside of mitosis of Myo1cKO embryos.**

Time lapse of a Myo1cKO embryo expressing mTmG labeling the plasma membrane (grey) and mCherry-EB3 (grey) labeling the mitotic spindle. Images were taken every 15 min. Scale bar, 10 *μ*m. White arrows pointing at fragments, red arrow pointing at the mitotic spindle.

**Movie 3: Mitotic spindle dynamics in control and Myo1cKO embryos.**

Time lapses of control (left) and Myo1cKO (right) embryo labeled with mTmG labeling the plasma membrane (grey) and mCherry-EB3 (grey) labeling the mitotic spindle. Images were taken every 15 min. Scale bar, 10 *μ*m. White arrows indicate example of disrupted spindle dynamics in Myo1c embryos.

**Movie 4: Cortical actin cap formation in proximity to DNA and fragment formation in Myo1cKO embryos.**

Time lapse of a Myo1cKO embryo expressing LifeAct-GFP (green) and labeled with SiRDNA (grey). Images were taken every 15 min. Scale bar, 10 *μ*m. White arrow indicates cotical actin cap, red arrow indicates fragment.

**Movie 5: Cortical Myh9 accumulation in proximity to DNA and fragment formation in Myo1cKO embryos.**

Time lapse of a Myo1cKO embryo expressing mTmG (magenta) and Myh9-GFP (green), labeled with SiRDNA (grey). Images were taken every 15 min. Scale bar, 10 *μ*m. White arrow indicates Myh9 accumulation, red arrow indicates fragment.

**Movie 6: Cortical Myh9 ring formation in proximity to DNA and fragment formation in Myo1cKO embryos.**

Time lapse of a Myo1cKO embryo expressing mTmG (magenta) and Myh9-GFP (green), labeled with SiRDNA (grey). Images were taken every 15 min. Scale bar, 10 *μ*m. White arrows indicate Myh9 accumulation in shape of ring, red arrow indicates fragment.

**Movie 7: Cortical Cdc42 activation in proximity to DNA in control and Myo1cKO embryos.**

Time lapses of control (left) and Myo1cKO (right) embryo labeled with SiRDNA (grey) and where one of the blastomere expresses a Cdc42 activity reporter (yellow). Images were taken every 15 min. Scale bar, 10 *μ*m. White arrow indicates Cdc42 activity accumulation. White line outlines the blastomere not expressing the Cdc42 reporter.

**Movie 8: Cortical actin accumulation in Myo1cKO embryos in presence and absence of Cdc42 activity.**

Time lapses of Myo1cKO embryos labeled with SiRDNA (grey) expressing LifeAct-GFP (green) and DNCdc42 (right) or not (left). Images were taken every 15 min. Scale bar, 10 *μ*m. White arrows indicate cortical region in close proximity to DNA.

**Movie 9: Cortical actin cap in GFP and Ect2-GFP expressing embryos.**

Time lapses of embryos labeled with SiRDNA (grey) and expressing LifeAct-RFP (green) and GFP (left, blue) or Ect2-GFP (right, blue). Images were taken every 15 min. Scale bar, 10 *μ*m. White arrows indicate cortical actin cap.

